# Spatial specificity of auxin responses coordinates wood formation

**DOI:** 10.1101/142885

**Authors:** Klaus Brackmann, Virginie Jouannet, Jiyan Qi, Theresa Schlamp, Karin Grünwald, Pablo Sanchez, Thomas Greb

## Abstract

Spatial organization of signaling events of the phytohormone auxin is fundamental for maintaining a dynamic transition from plant stem cells to differentiated descendants. The cambium, the stem cell niche mediating wood formation, fundamentally depends on auxin signaling but its exact role and spatial organization is obscure. Here, we show that, while auxin signaling levels increase in differentiating cambium descendants, a moderate level of signaling in cambial stem cells is essential for cambium activity. We identify the auxin-dependent transcription factor ARF5/MONOPTEROS to cell-autonomously restrict the number of stem cells by attenuating the activity of the stem cell promoting *WOX4* gene. In contrast, ARF3 and ARF4 function as cambium activators in a redundant fashion from outside of WOX4-expressing cells. Our results reveal an influence of auxin signaling on distinct cambium features by specific signaling components and allow the conceptual integration of plant stem cell systems with distinct anatomies.

## Introduction

In multicellular organisms, communication between cells is essential for coordinated growth and determination of cell fate. In plants in particular, the flexible regulation of cellular properties by hormone signaling is important throughout the whole life cycle. This is because plants are sessile and continuously adapt their growth and development to their local environment. The basis of this plastic growth mode are local stem cell niches at the tips and along plant growth axes, called meristems^1^. The tip-localized shoot and root apical meristems (SAM & RAM) are essential for primary, or longitudinal, growth of shoots and roots, respectively. In turn, the vascular cambium is the predominant lateral meristem forming a cylinder of indeterminate stem cells at the periphery of growth axes and mediating radial growth by producing the vascular tissues phloem and xylem in a bidirectional manner^2, 3^. This production is the basis of wood formation and is, thus, essential for the accumulation of a large proportion of terrestrial biomass.

The plant hormone auxin plays pivotal roles in local patterning and maintenance of stem cell niches in the SAM and RAM. In the SAM, auxin signaling is low in stem cells and increases during recruitment of cells for organ formation^4, 5, 6^. Cell wall modulation and the formation of vascular strands are two aspects promoted by auxin in this context^7, 8^. In contrast, a maximum of auxin signaling is present in the quiescent center (QC) and the surrounding stem cells in the RAM and cell differentiation is, at least partly, driven by a decrease in signaling levels^9, 10^. Therefore, the functions of auxin in both meristems are different and adapted to distinct niche requirements.

For the cambium, the role of differential auxin signaling along the radial sequence of tissues is still obscure. In *Arabidopsis* stems apex-derived auxin is transported basipetally and distributed laterally across the cambial zone by the auxin exporters PIN-FORMED1 (PIN1), PIN3, PIN4 and PIN7^11, 12^. Indeed, direct auxin measurements in *Populus* and *Pinus* trees showed that the concentration of the major endogenous auxin indole-3-acetic acid (IAA) peaks in the center of the cambial zone and gradually declines towards differentiating xylem and phloem cells^13, 14, 15^. This observation prompted the idea that, in analogy to the situation in the RAM, radial auxin concentration gradients contribute to the transition of cambium stem cells to secondary vascular tissues^16, 17^. Consistently, ubiquitous repression of auxin responses by expressing a stabilized version of the auxin response inhibitor PttIAA3 reduces the number of cell divisions in the cambium region of hybrid aspen trees^18^. In addition, however, the zone of anticlinal cell divisions characteristic for cambial stem cells is enlarged in PttIAA3 overexpressing trees. This suggests that auxin signaling not only promotes cambium proliferation but also spatially restricts stem-cell characteristics within the cambium area^18, 19^. Indeed, especially xylem formation is associated with a local increase of auxin signaling in other contexts^10, 20, 21, 22^ which supports a role of auxin in the recruitment of cells for differentiation similarly as in the SAM. Therefore, it is currently unclear whether auxin signaling is predominantly associated with stem cell-like features or cell differentiation in the context of radial plant growth or how a positive effect on cambium proliferation and on the differentiation of vascular tissues is coordinated.

As a central cambium regulator, the WUSCHEL-RELATED HOMEOBOX4 (WOX4) transcription factor imparts auxin responsiveness to the cambium^23^. Equivalent to the role of WUSCHEL (WUS) and WOX5 in the SAM and RAM^24, 25^, WOX4 activity maintains stem cell fate^23, 26^. In turn, *WOX4* transcription is stimulated by the leucine-rich repeat receptor-like kinase (LRR-RLK) PHLOEM INTERCALATED WITH XYLEM (PXY). Importantly, the expression domains of the *WOX4* and *PXY* genes presumably overlap and are considered to mark cambium stem cells^23, 26, 27, 28^. However, a bipartite organization of the cambium zone was shown recently with *PXY* being expressed only in the xylem-facing part^29^. Whether this organization reflects the existence of two distinct stem cell pools feeding xylem and phloem production, respectively, has still to be determined.

Here, we identify functional sites of auxin signaling in the *Arabidopsis* cambium by local short-term modulation of auxin biosynthesis and signaling. We reveal that, while cambial stem cells do not appear to be a site of elevated auxin signaling, auxin signaling in these cells is required for cambium activity. By analyzing transcriptional reporters and mutants of vasculature-associated AUXIN RESPONSE FACTORs (ARFs), we identify ARF3, ARF4 and ARF5 as cambium regulators with different tissue-specificities as well as distinct roles in cambium regulation. Remarkably, whereas ARF3 and ARF4 act redundantly as more general cambium promoters, ARF5 acts specifically in cambium stem cells. In depth analysis of the auxin- and ARF5-dependent transcriptome in those cells, together with genetic analyses, indicates that the ARF5-dependent repression of *WOX4* is an essential aspect of auxin signaling during cambium regulation.

## Material and Methods

### Plant Material

All plant lines used in this study were *Arabidopsis thaliana* (L.) Heynh. plants of the accession Columbia (Col-0), except for the *mp-B4149* mutant, which has the Utrecht background^30, 31^. The *arf3-1* (SAIL_1211_F06, N878509), *ett-13* (SALK_040513, N540513), *mp-S319* (SALK_021319, N521319), *wox4-1* (GK_462GO1, N376572) and *pxy-4* (SALK_009542, N800038) mutants, as well as the *pDR5rev:GFP* reporter line (N9361^32^), were ordered from the Nottingham Arabidopsis Stock Centre (NASC). The *mp-B4149* and *arf4-2* (SALK_070506) mutant were provided by Dolf Weijers (University of Wageningen, The Netherlands) and Alexis Maizel (COS Heidelberg, Germany), respectively. Genotyping was performed by PCR using primers listed in Table S1. Genotyping of *mp-B4149* was done as described previously^33^ with the modification of using the MP_for8/MP_rev8 primer pair for amplification.

### Plant Growth and Histological Analyses

Plants destined for histology were grown and analyzed as described previously^23, 28, 34^. In brief after 3 weeks of growth in short day (SD) conditions (8 h light and 16 h dark) plants were transferred to long day (LD) conditions (16 h light and 8 h dark) to induce flowering and used for histology at a height of 15-20 cm. Stem segments of at least 1 cm in length (incl. the stem base) were harvested, embedded in paraffin and sectioned using a microtome (10 μm sections). After deparaffinization, sections were stained with 0.05% toluidine blue (Applichem), fixed with Entellan (Merck) and imaged using a Pannoramic SCAN digital slide scanner (3DHistech). Pictures were analyzed as described previously^34^ using Pannoramic Viewer 1.15.4 software (3DHistech). For quantitative analyzes at least five plants were analyzed for each data point.

### Statistical Analyses

Statistical analyses were performed using IBM SPSS Statistics for Windows, Version 21.0. Armonk, NY: IBM Corp. Means were calculated from measurements with sample sizes as indicated in the respective figure legends. In general, all displayed data represents at least two independent, technical repetitions, unlike otherwise indicated. Error bars represent ± standard deviation. All analysed datasets were prior tested for homogeneity of variances by the Levene statistic. Significant differences between two datasets were calculated by applying a Welch’s t-tests or Student’s t-test depending on the homogeneity of variances. The significance thresholds were set to p-value < 0.05 (indicated by one asterisk). For multiple comparisons between three or more datasets, a One-way ANOVA was performed, using a confidence interval (CI) of 95% and a post-hoc Bonferroni for comparisons of data sets of homogenous variances or a post-hoc Tamhane-T2 in case variances were not homogenous.

### Sterile culture

Adventitious root formation in the strong *arf5* mutant allele *mp-B4149* was induced with some minor modifications as described previously^35, 36^. Seeds were liquid sterilized by 70% ethanol and incubation in 5% sodiumhyperchloride followed by three washes with ddH2O. After stratification at 4°C in the dark for 3 days, seeds were sown on ½ Murashige-Skoog (MS) medium plates (incl. B5 vitamins) in rows and grown vertically. After 7 days of growth under short-day (SD; 8 h light, 16 h dark) conditions, rootless mutant as well as wild type looking seedlings from the segregating population were bisected with a scalpel as described previously^36^ and transferred to adventitious root inducing medium (½ MS (incl. B5 Vitamins) + 1.5 *%* sucrose + 3 μg/ml indole butyric acid + 0.7 % agar + 50μg/ml ampicillin^35^). After additional two weeks of growth under SD conditions, successfully rooted seedlings were transferred to soil and grown under SD conditions for one additional week before they were transferred to LD conditions. Plants that survived the transfer, were genotyped for *mp-B4149* and only wild type and homozygous mutant plants were used for histological analysis at a plant height of 15-20 cm as described in the previous section.

### Plasmid construction

The *pPXY:CFP* and *pWOX4:YFP* reporters were described previously^23, 28^. To avoid diffusion, all fluorescent proteins were targeted to the endoplasmatic reticulum (ER) by fusing them to the corresponding sequence motif (ER + HDEL motif^37^). For generating *pDR5revV2:YFP (pKB46),* we initially inserted the *ADAPTOR PROTEIN-4 MU-ADAPTIN (AP4M,* At4g24550) terminator, amplified from genomic DNA using the At4g24550_for1/At4g24550_rev1 primer pair, into *pLC075* containing the *DR5revV2* promoter fragment^38^ using BamHI/XhoI restriction sites. A fragment carrying the *ER-EYFP-HDEL* coding sequence (CDS) was inserted in the resulting *pLC075:AP4Mterm* using the BamHI restriction site, to obtain *pLC075:YFP:AP4Mterm.* The complete reporter fragment was inserted in the binary vector *pGreenII017^39^* using KpnI/XhoI restriction sites. For generating *p35S:Myc-GR-bdl (pKB9),* the *Myc-GR-bdl* fragment was amplified from genomic DNA of *pRPS5a:Myc-GR-bdl*^40^ using the Myc_for1/BDL_rev3 primer pair. The resulting fragment was inserted in the *pGreen0229* vector^39^ containing the 35S promoter (*pGreen0229-35S*) using XbaI/BamHI restriction sites. To produce *pAlcA:iaaM(pKB2)* the *iaaM* CDS was amplified from *piaaM (pIND:IND-iaaM)^41^* using the IAAMfor1/IAAMrev1 primer pair and introduced into *pGreen0229-AlcA^42^* using AatII/EcoRI restriction sites. *pWOX4:AlcR (pTOM55*) was produced by amplifying the *WOX4* promoter using the primers WOX4for11/WOX4ref9 and inserting the resulting fragment into *pAlcR-GUS^42^* using SpeI/NotI sites. The *pPXY:Myc-GR-bdl (pKB45*) construct was generated by cloning the *Myc-GR-bdl* fragment, amplified from *pKB9* using the Myc_for5/BDL_rev7 primer pair, in *pGreen0229* containing the PXY promoter (*pTOM50*^28^) using NcoI/Cfr9I restriction sites. The promoter regions of BDL^43^ were amplified from genomic DNA using the BDL_for2/BDL_rev4 & BDL_for3/BDLrev5 primer pairs. Both fragments were cloned into *pGreenII0179* using NotI/XbaI & Cfr9I/KpnI restriction sites, respectively. The resulting plasmid (*pKB27*) was used to produce the *pBDL:YFP (pKB28* using ER-EYFP-HDEL) and *pBDL:Myc-GR-bdl (pKB29*) constructs by inserting fragments carrying the respective CDSs using NcoI/Cfr9I restriction sites. For generating *ARF3*, *ARF4* and *ARF5* reporter constructs, promoter regions of the three genes were amplified from genomic DNA using the ARF3for1/ARF3rev1 & ARF3for2/ARF3rev2, ARF4for1/ARF4rev1 & ARF4for2/ARF4rev2 and MP_for7/MP_rev5 & MP_for5/MP_rev6 primer pairs. Both fragments were cloned for each gene into *pGreen0229* using NotI/BamHI & BamHI/KpnI (*ARF3),* NotI/SpeI & Cfr9I/KpnI (ARF4) and NotI/BamHI & BamHI/ApaI (*ARF5*) restriction sites. The resulting plasmids (*pKG40 (ARF3), pKG41 (ARF4*) & *pKB3* (ARF5)) were used to produce the *pARF3:YFP (pKB30), pARF4:YFP (pKB31), pARF5:YFP (pKB24*) and *pARF5:mCherry (pKB4*) constructs by inserting fragments carrying the respective CDSs using BamHI, SpeI and NdeI/XhoI restriction sites, respectively. To produce *p35S:GR-ARF3 (pKB42*) and *p35S:GR-ARF5 (pKB17*) we amplified the GR open reading frame from *pKB9* using the GR_for1/GR_rev3 primer pair and inserted the resulting fragment in *pGreen0229-35S* using XbaI/Cfr9I restriction sites. An additional unannotated SalI restriction site in the *pGreen0229* backbone was removed by PCR-based silent mutagenesis using the NOS-mut_for1/NOS_mut_rev1 primer pair. In the resulting *pGreen0229-35S:GR (pKB41*) vector we inserted the *ARF3* and *ARF5* CDS, amplified from cDNA using the ARF3_for4/ARF3_rev4 and MP_for16/MP_rev14 primer pair, using SalI/Cfr9I and SalI/EcoRI restriction sites, respectively. *pPXY:GR-ARF3 (pKB43*) was generated by cloning the *GR-ARF3* fragment, amplified from *pKB42* using the GR_for5/ARF3_rev4 primer pair, in *pTOM50* using NcoI/Cfr9I restriction sites. For generating *pPXY:GR-ARF5ΔIII/IV (pKB25),* the *GR-ARF5ΔIII/IV* fragment with a stop codon was amplified from *pKB17* using the MP_for18/MP_rev16 primer pair and inserted in *pTOM50* using NcoI/Cfr9I restriction sites. For generating the *pWOX4:LUC (firefly);p35S:LUC (Renilla) (pKB55*) reporter the *pZm3918:LUC (firefly*) fragment in *pZm3918:LUC (firefly);p35S:LUC (Renilla) (pGreen-LUC-REN*) was excised by digest with KpnI/XbaI and replaced by a *pWOX4:LUC* (firefly) fragment previously excised from *pWOX4:LUC (pMS80*) using KpnI/NheI restriction sites. To produce *p35S:ARF3 (pKB44*) the *ARF3* CDS was amplified from *pKB42* using the ARF3_for6/ARF3_rev4 primer pair and introduced it in *pGreen0229-35S* using XbaI/Cfr9I restriction sites. For generating *p35S:ARF5ΔIII/IV (pKB40*) the ARF5ΔIII/IV CDS was amplified from *pKB25* using MP_for17/MPrev15 and introduced in *pGreen0229-35S* using XbaI/EcoRI restriction sites. All constructs were sequenced and after plant transformation by floral dip^44^, single copy transgenic lines were identified by Southern blot analyses and representative lines used for crosses and further analyses. All primers mentioned in this section are listed in Table S1.

### Confocal microscopy

For imaging fluorescent reporter lines in the stem rough hand sections were taken with a razor blade (Wilkinson Sword) and analyzed using an LSM 780 spectral confocal microscope (Carl Zeiss) equipped with the Zen 2012 software (Carl Zeiss). Stem sections (except for *pARF5:mCherry*) were counterstained for 5 min with 5 μg/ml propidium iodide (PI; Merck) dissolved in tap water. PI was excited at 561 nm (DPSS laser) and detected at 590-690 nm. YFP was analysed with excitation at 514 nm (Argon laser) and detection at 516-539 nm. CFP was excited at 458 nm (Argon laser) and detected at 462-490 nm, while GFP was excited at 488 nm (Argon laser) and detected at 499-544 nm. mCherry reporter activity was analyzed with excitation at 561 nm (DPSS laser) and detection at 597-620 nm. Transmitted light pictures were generated using the transmission photo multiplier detector (T-PMT) of the microscope. 5-day-old *Arabidopsis* seedlings were counterstained with the cell membrane dye FM^®^ 4-64 (Thermo Fisher Scientific) to visualize cell borders as described previously^45^. FM^®^ 4-64 was excited at 561 nm (DPSS laser) and detected at 653-740 nm.

### Pharmacological treatments

Stock solutions of 25 mM Dex (VWR) dissolved in 100% Ethanol and 10 mM cycloheximide (Cyclo; Carl Roth) dissolved in ddH2O were freshly prepared prior to use. For long-term Dex-treatments, plants were initially grown for three weeks without treatment in SD conditions to circumvent growth defects during early plant development. Plants were then transferred to LD conditions and watered twice a week with either 15 μM Dex (25 mM Dex stock diluted in tap water) or mock solution (equal amount of 100 % Ethanol in tap water) until they reached a height of 15-20 cm and were harvested for histology. For short-term Dex-treatments 15-20 cm tall plants were dipped headfirst in 15 μM Dex (25 mM Dex stock diluted in tap water + 0.02% Silwet) or Mock solution (equal amount of 100 *%* Ethanol in tap water + 0.02% Silwet) for 30 sec. Subsequently, plants were transferred to LD growth conditions, watered with 15 μM Dex or Mock solution and incubated until harvest of second internodes for RNA isolation. For additional short-term Cyclo treatment, 10 mM Cyclo stock was added to the 15 μM Dex and Mock solution to a final concentration of 10 μM and the plants were treated in the same way as described before. For inducing the AlcA/AlcR system^42^, plants where grown for three weeks in SD, transferred to long day until bolting. When plants where 0.5 – 3 cm tall they were put under a plastic dome together with 2 x 15 ml 70% ethanol in round petri dishes and left overnight. Plants where harvested 10 days after induction and wild type plants were around 25 cm tall.

### Transient reporter activity assays

Transient activity assays were performed as previously described^46^. In brief protoplasts derived from an *Arabidopsis* (Col-0) dark-grown root cell suspension culture (kindly provided by Claudia Jonak) were isolated and transfected as previously described^47^. For transfection, we used 10 μg of reporter construct (*pKB55*) containing *p35S:LUC (Renilla*) as an internal control and 10 μg of each effector construct. The transfected protoplasts were diluted with 240 mM CaCl2 (1:3) followed by cell lysis and dual-luciferase assay using the Dual-Luciferase Reporter Assay System (Promega) and following the manufacturer’s instructions. Luminescence was measured using a Synergy H4 Hybrid Multiplate Reader (BioTek). For each reporter/effector combination 3-5 technical replicates were done and the experiments repeated at least three times. For experimental analysis Firefly Luciferase activity was normalized to Renilla Luciferase activity.

### RNA Preparation and qRT-PCR

Frozen plant material from second internodes or the stem base (incl. 5 mm above) of 15-20 cm tall plants (three biological replicates (three plants each) per genotype/treatment) were pulverized with pestle and mortar and RNA was isolated by phenol/chlorophorm extraction as described previously^48^ with the modification of two additional concluding 70% EtOH washes. RNA elution in RNase-free water was followed by treatment with RNase-free DNase (Thermo Fisher Scientific) and reverse transcription (RevertAid First Strand cDNA Synthesis Kit; Thermo Fisher Scientific). cDNA was diluted 1:25 prior to amplification. qRT-PCR was performed using SensiMix™ SYBR^®^ Green (Bioline Reagents Ltd) mastermix and gene specific primers (listed in Table S1), in a Roche Lightcycler480 following the manufacturer’s instructions. Experiments were performed in triplicates with plant material of three plants being pooled for each replicate. Two reference genes (ACT2 and EIF4a) were used to normalize our signal. Error bars: ± standard deviation. Raw amplification data were exported and further analysis and statistical tests were done using Microsoft Excel 2010.

### Transcriptional Profiling

10 μg of total RNA for each sample were treated with RNase-free DNase (Thermo Fisher Scientific) and purified using RNA-MiniElute columns (Qiagen) following the manufacturer’s protocol. Library preparation and next-generation-sequencing (NGS) was performed at the Campus Science Support Facilities (CSF) NGS Unit (www.csf.ac.at) using HiSeqV4 (Illumina) with single end 50-nucleotide reads. Reads were aligned to the *Arabidopsis thaliana* Columbia (TAIR10) genome using CLC Genomics Workbench v7.0.3 and analyzed using the DESeq package from the R/Bioconductor software^49^. Dex-treated samples were compared to mock-treated samples with a stringency of p-value < 0.05. Data processing was further analyzed using VirtualPlant 1.3^50^ Gene Sect and BioMaps with a cut-off p-value < 0.05 and cut-off p-value <0. 01, respectively. Data was aligned to The *Arabidopsis* Information Resource (TAIR)-databases (77) and as background population for all analysis the *Arabidopsis thaliana* Columbia (TAIR10) genome was used. Further data processing was done in Microsoft Excel 2010.

### Accessing RNA sequencing data

Raw data produced in this study have been uploaded to NCBI’s Gene Expression Omnibus (GEO) database^51, 52^ and are accessible through GEO Series accession number GSE98193.

## Results

### Local auxin responses in stem cells stimulate cambium activity

In *Arabidopsis* stems, the activity of the common auxin response marker *pDR5rev:GFP*^32^ was detected in vascular tissues (phloem and xylem) and cortical cells prior and during cambium initiation (Figures 1A, S1A)^23^. However, there was no overlap with *pWOX4:YFP*^23^ or *pPXY:CFP^28^* reporter activities, the two canonical markers for cambium stem cells (Figure 1A-C, S1A-C)^23^. This suggested that auxin signaling in those cells occurs at low levels or is even absent. To decide between both possibilities, we generated a plant line expressing an endoplasmatic reticulum (ER)-targeted Yellow Fluorescent Protein (YFP) under the control of the high affinity *DR5revV2* promoter which recapitulated the pattern of *DR5revV2* activity previously reported in roots (Figures S1D-F)^38^. In the second internode of elongated shoots, *pDR5rev:GFP* and *pDR5revV2:YFP* activities were congruent but *pDR5revV2:YFP* activity also included the whole cortex as well as cambium cells marked by *pPXY:CFP* activity (Figure S1G-I). Immediately above the uppermost rosette leaf (denoted as stem base throughout the text), stem anatomy shows a secondary configuration, which is characterized by a continuous domain of cambium activity^23^. At this position, the expression domain of *pDR5revV2:YFP* was again broader than the domain of *pDR5rev:GFP* activity substantially overlapping with *pPXY:CFP* activity (Figure 1D-F). Based on these observations, we concluded that the auxin signaling machinery is active in PXY-positive cambial stem cells.

**Figure 1:**
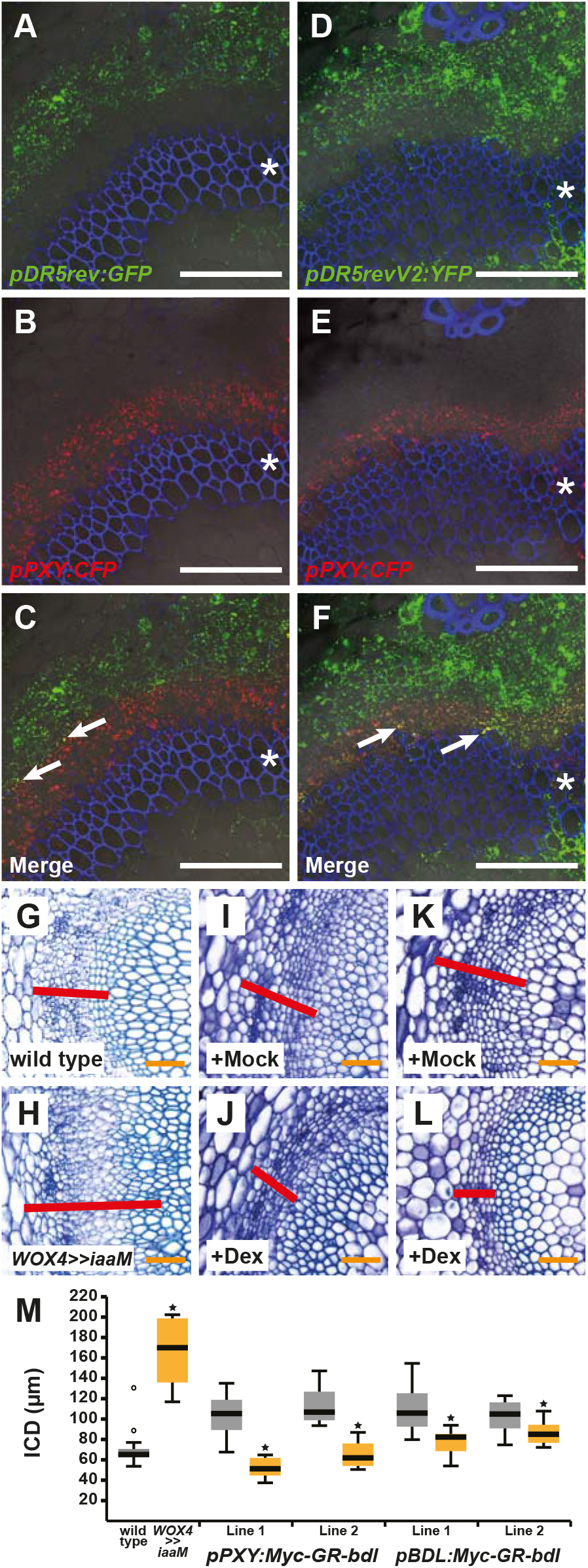
**A-F:** Confocal analyses of stem bases from plants containing the auxin response markers *pDR5rev:GFP* (A-C) or *pDR5revV2:YFP* (D-F) and the stem cell marker *pPXY:CFP.* Overlapping foci between *pPXY:CFP* (red) and the respective auxin response marker activities are marked by arrows (C & F). Asterisks mark the vascular bundles. Size bars represent 100 μm. Propidium iodide (PI) staining in blue. **G-L:** Toluidine blue stained cross sections at the stem base of wild type (G) *pWOX4:AlcR; pAlcA:iaaM* (H), *pPXY:-Myc-GR-bdl* (I, J) and *pBDL:Myc-GR-bdl* (K, L) plants after long-term EtOH (G, H), mock (I, K) or dexamethasone (J, L) treatment. Interfascicular regions are shown and interfascicular cambium-derived tissues (ICD) are marked (red bar). Size bars represent 50 μm. **M:** Quantification of ICD tissue extension at the stem base of wild type, *pWOX4:AlcR; palcA:iaaM (WOX4>>iaaM), pPXY:Myc-GR-bdl* and *pBDL:Myc-GR-bdl* plants after long-term EtOH (wild type & *WOX4>>iaaM),* mock (grey) or dexamethasone (yellow) treatment. Homogeneity of variance was tested by F-test and accordingly Student’s T-test (*pPXY:Myc-GR-bdl* (line 2) p=9.82E-06 & *pBDL:Myc-GR-bdl* p=0.001 & p=0.03) or Welch’s T-test (wild type and *WOX4>>iaaM* p=9.24E-09 & *pPXY:Myc-GR-bdl* (line 1) p=5.2E-06) were performed comparing wild type and *WOX4>>iaaM* and mock and Dex, respectively (Sample sizes n=8-16). Significance is indicated by asterisks.

To see whether cambium activity was positively correlated with auxin levels in PXY-positive cells, we used the *WOX4* promoter, whose activity fully recapitulated the *PXY* promoter activity (Figure S2A-I), for expressing a bacterial tryptophan monooxygenase (iaaM) in an inducible manner^42^. iaaM converts endogenous tryptophan to the IAA precursor indole-3-acetamide and was used before to boost endogenous IAA levels in *Arabidopsis*^53^. As a read out for cambium activity, we determined the amount of interfascicular cambium-derived (ICD) tissues^34^. Indeed, ethanol-based iaaM induction substantially stimulated the production of ICD tissues (Figure 1G, H, M, Figure S2J-N) demonstrating that an increase of auxin biosynthesis in PXY-positive stem cells stimulates cambium activity.

To determine to which extent downstream components of the auxin signaling cascade are required in those cells, we blocked ARF activity by expressing a dexamethasone (Dex)-inducible variant of the stabilized AUX/IAA protein BODENLOS (Myc-GR-bdl)^40, 54^ under the control of the *PXY* promoter. Consistent with a role of ARF activity in cambium regulation, Dex-treatments of *pPXY:Myc-GR-bdl* plants resulted in a strongly reduced amount of ICD tissues at the stem base (Figure 1I, J, M) but not in an altered overall growth habit (Figure S2O). Strikingly, Dex-treated *pPXY:Myc-GR-bdl* plants showed an even more pronounced repression of IC activity than the inhibition using the *BDL* promoter^43^ (Figure 1K, L, M, Figure S2P) whose activity was very broad including also PXY-positive cells (Figure S3A-F). These observations indicated that, local auxin signaling in PXY-positive stem cells stimulates cambium activity.

### *ARF3, ARF4* and *ARF5* genes are expressed in cambium associated cells

To identify ARFs active in cambium stem cells, we mined public transcriptome datasets and found the *ARF3/ETTIN*, *ARF4* and *ARF5/MONOPTEROS* genes to be co-induced with *WOX4* and *PXY* during cambium initiation^23^. Indeed, *pARF3:YFP, pARF4:YFP, pARF5:YFP* promoter reporters were active in cambium-related cells at the stem base and the second internode (Figure 2A, D, G, Figure S4A, D, G). However, while *pARF3:YFP* and *pARF4:YFP* reporters were active in rather broad domains including the phloem, the xylem and, partly, pPXY:CFP-positive cells (Figure 2A-F, Figure S4A-F), *pARF5:YFP* was exclusively active in cells marked by *pPXY:CFP* activity (Figure 2G-I, Figure S4G-I). Moreover, in second internodes, *pARF3:YFP* and *pARF4:YFP* activities were both detected in the starch sheath, the innermost cortical cell layer which is considered to serve as the origin of the IC (Figure S4C & F arrows)^55, 56^ while *pARF5:YFP* activity was restricted to vascular bundles (Figure S4G-I). Indicating also a temporal difference between *ARF3/4* and *ARF5* activities, *pARF3:YFP* and *pARF4:YFP* reporters were active together with *pPXY:CFP* in interfascicular regions at positions approximately 5 mm above the stem base (Figure S5A-F) where cortical cells start dividing to form the IC^34^. In contrast, no *pARF5:YFP* activity was detected in the same cortical cells (Figure 2J-L). This observation suggested that *ARF5* expression follows the expression of *PXY* during cambium initiation and is not active during early steps of cambium initiation. Consistently, in *pxy-4* mutants where IC formation is largely absent in stems (Figure 5E; ^57^) a *pARF5:mCherry* promoter reporter was only active in vascular bundles but not in interfascicular regions (Figure S5G-I). Taken together, these observations were in line with a role of *ARF3* and *ARF4* as promoters of cambium activity and a role of *ARF5* as a modulator of the established cambium.

**Figure 2:**
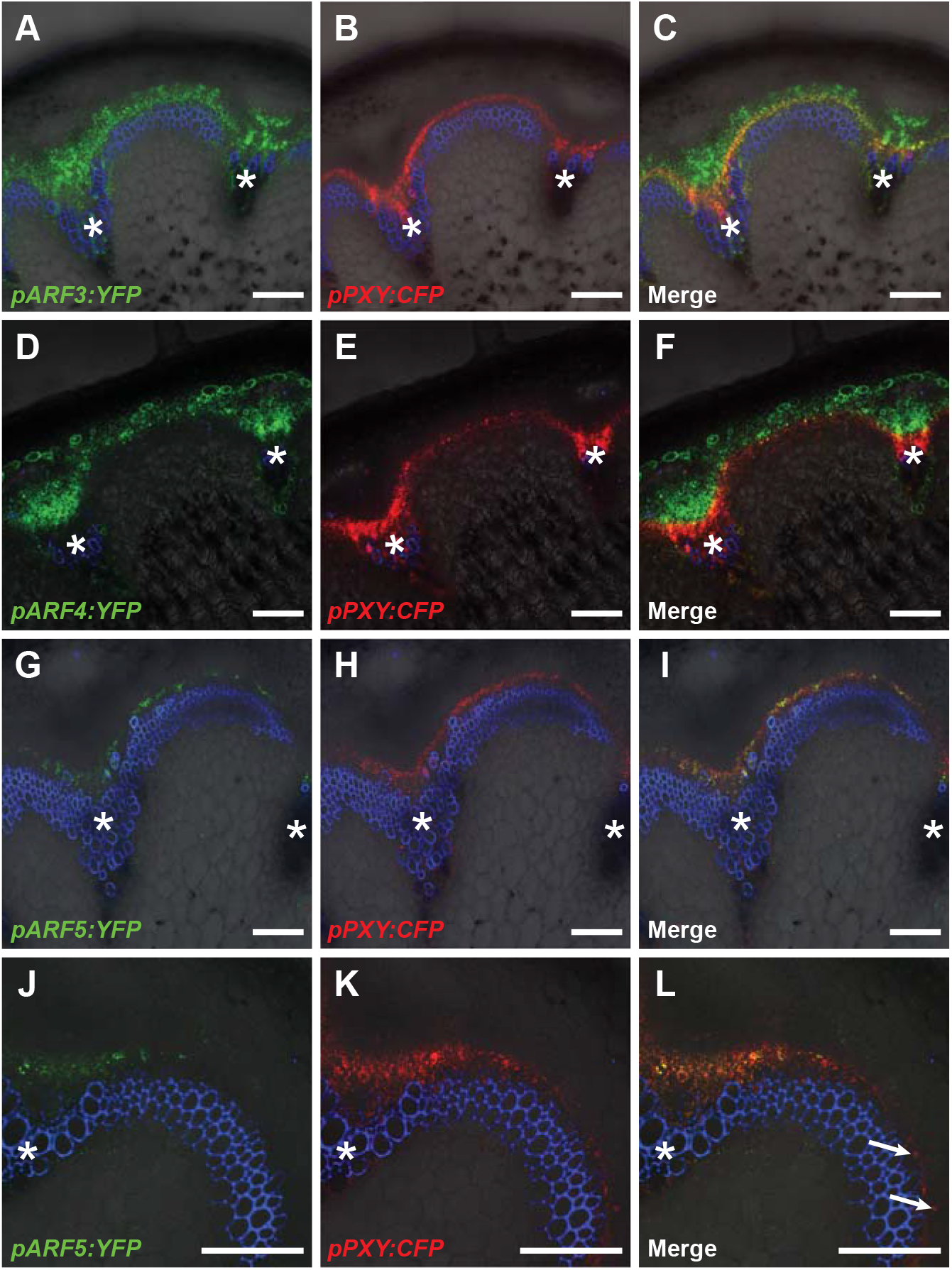
**A-I:** Confocal analyses of stem bases of plants containing *pARF3:YFP* (A-C), *pARF4:YFP* (D-F) and *pARF5:YFP* (G-I), respectively, and the stem cell marker *pPXY:CFP.* Asterisks mark the vascular bundles. Size bars represent 100 μm. **J-L:** Confocal analyses of cross sections from 5 mm above the stem base (transition zone) of plants containing *pARF5:YFP* and the stem cell marker *pPXY:CFP.* Arrows mark cells in the interfascicular region displaying CFP (red) but no YFP (green) activity. Size bars represent 100 μm. PI staining in blue.

### ARF control of cambium proliferation

To find indications for these roles, we analyzed cambium activity in mutants for the respective *ARF* genes (Figure S5J-L). Consistent with a positive effect of *ARF3,* both weak and strong *arf3* mutants^58, 59^ showed significantly reduced cambium activity (Figure 3A, B, E, F, M). This reduction was further increased upon depletion of *ARF4* activity by introducing the *arf4-2* mutation^59^ into the respective *arf3* mutant backgrounds (Figure 3C, D, G, H, M). Consequently, we concluded that cambium activity is positively regulated by *ARF3* and *ARF4*, which, as in other contexts^59, 60^, act in a concerted fashion.

**Figure 3:**
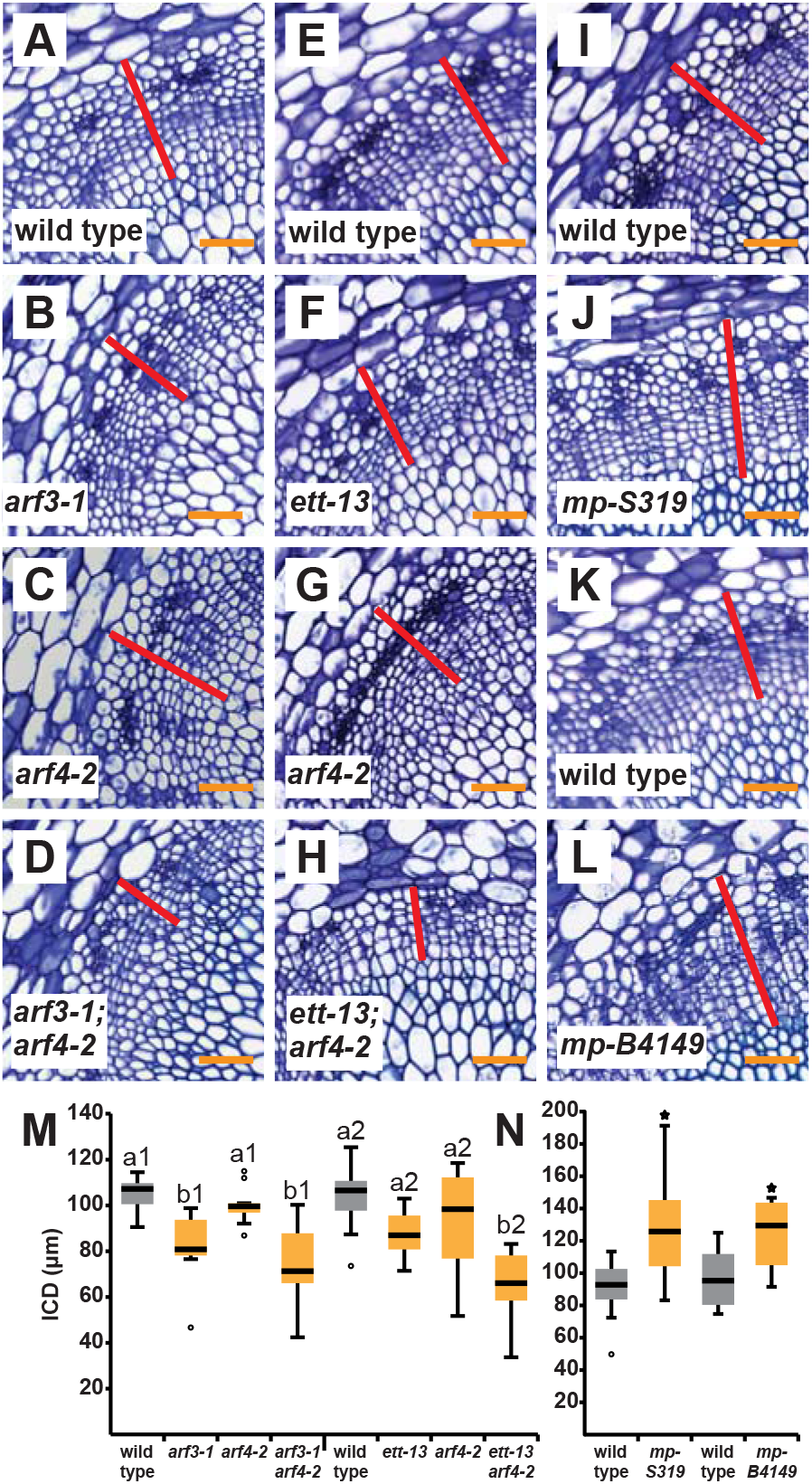
**A-L:** Toluidine blue stained cross sections at the stem base of wild type and arf single and multiple mutant plants. Genotypes are indicated. Interfascicular regions are shown and interfascicular cambium-derived tissues (ICD) are marked (red bar). Size bars represent 50 μm. **M, N:** Quantification of ICD tissues at the stem base of wild type, *arf3/4* single and double mutants (M) and *arf5/mp* mutant plants (N). Statistical groups indicated by letters were determined by one-way ANOVA with post hoc Bonferroni (CI 95%; Sample size n=8-10) (M). Student’s T-test was performed comparing wild type and *mp-S319* (p=0.003) and wild type and *mp-B4149* (p=0.03), respectively (Sample sizes n=4-12) (N). Significance is indicated by asterisks.

In contrast, cambium activity was enhanced in the hypomorphic *arf5* mutant *mp-S319^61^* (Figure 3I, J, N) suggesting that *ARF5* counteracts cambium proliferation. To confirm this role, we generated adult plants of the strong *ARF5* loss-of-function mutant *mp-B4149* ^30^ and wild type plants through tissue culture^35, 36^. As before, *mp-B4149* plants showed enhanced ICD formation comparable to *mp-S319* mutants (Figure 3K, L, N). Further confirming a negative effect of ARF5 on cambium activity, ubiquitous expression of a Dex-dependent GR-ARF5 protein fusion using the *35S* promoter^62^ led to significantly reduced tissue production under long-term induction (Figure 4A-C, J).

**Figure 4:**
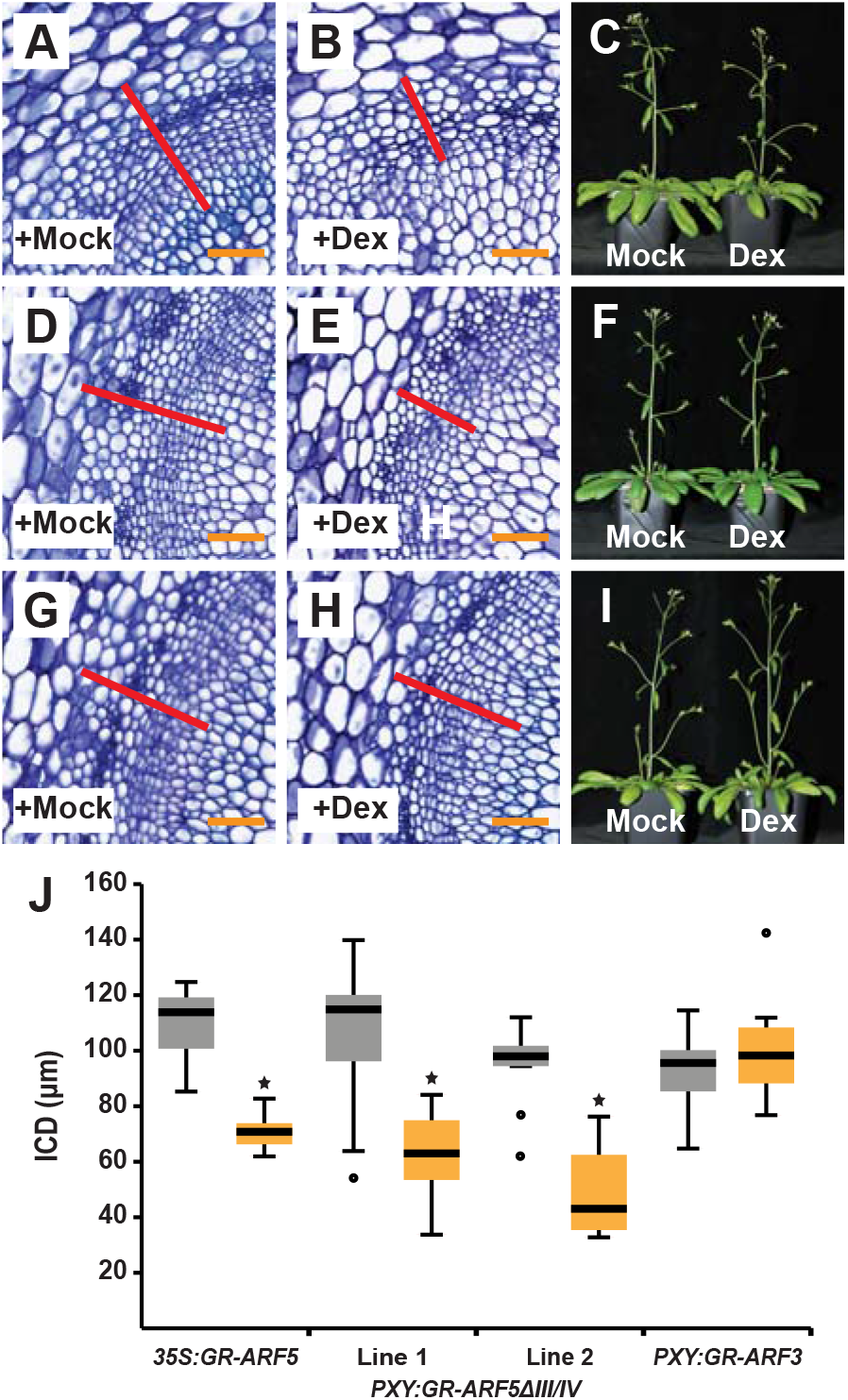
**A-B, D-E, F-G:** Toluidine blue stained cross sections at the stem base of *p35S:GR-ARF5* (A-B), *pPXY:GR-ARF5Δ III/IV* (D-E) and *pPXY:GR-ARF3* (G-H) plants after long-term mock (A, D & G) or dexamethasone (B, E & H) treatment. Interfascicular regions are shown and ICD tissues are marked (red bar). Size bars represent 50 μm. **C, F, I:** Overview pictures of *p35S:GR-ARF5* (C), *pPXY:GR-ARF5ΔIII/IV* (F), *pPXY:GR-ARF3* (I) plants after long-term mock or Dex treatment. **J:** Quantification of ICD tissue extension at the stem base of *p35S:GR-ARF5, pPXY:GR-ARF5ΔIII/IV* and *pPXY:GR-ARF3* plants after long-term mock (grey) or dexamethasone (yellow) treatment. Student’s T-test (*p35S:GR-ARF5* p=3.03E-05, *pPXY:GR-ARF5ΔIII/IV* (line 2) p=1.34E-05, *pPXY:GR-ARF3* p=0.37) and Welch’s T-test (*pPXY:GR-ARF5ΔIII/IV* (line 1) p=9.72E-04) were performed comparing mock and Dex treatment (Sample sizes n=6-10). Significance is indicated by the asterisk.

To test whether the identified ARFs function in PXY-positive stem cells, we first employed the *PXY* promoter to express GR-ARF5ΔIII/IV, a truncated variant of ARF5 lacking the domains III and IV releasing it from AUX/IAA-based repression^63^. Indeed, long-term Dex treatment of *pPXY:GR-ARF5ΔIII/IV* plants resulted in reduced cambium proliferation (Figure 4D-F, J) arguing for a stem cell-specific role of ARF5. In contrast, the same treatment of a *pPXY:GR-ARF3* line, did not influence cambium activity (Figure 4G-I, J, see below) arguing against a rate-limiting role of the non-AUX/IAA-dependent^64^ ARF3 protein in those cells. Collectively, we concluded that ARF3 and ARF4 on one side and ARF5 on the other side represent two subgroups of ARF transcription factors with differences in both their spatio-temporal expression and roles in cambium regulation.

### ARF5 restricts the number of undifferentiated cambium cells

To dissect the ARF5-dependent control of cambium stem cells, we took advantage of the DEX-inducibility of our *pPXY:GR-ARF5ΔIII/IV* and of a *p35S:Myc-GR-bdl* line. By determining transcript abundance of the direct ARF5 targets *ATHB8* and *PIN1^65,^* ^66^ at different time points after Dex treatment, 3 h of treatment was identified as being optimal for observing short-term effects on gene activity (Figure S6A, B). After establishing genome-wide transcript profiles at that time point, we identified a common group of 600 genes with altered transcript levels in both the *pPXY:GR-ARF5ΔIII/IV* and the *p35S:Myc-GR-bdl* line (p < 0.01; Figure 5A & Table S2). The 600 genes represented various functional categories including primary auxin response (IAAs, SAURs, GH3s), xylem and phloem formation (IAA20 & IAA30^22^, REV^12^, CVP2 & CVL1^67^) and cell wall modifications (PMEs & EXPs) (Figure S6C & Table S3). Moreover, the 312 genes that were induced by *pPXY:GR-ARF5ΔIII/IV* and repressed by *p35S:Myc-GR-bdl* (Figure 5A; Table S2), overlapped significantly with a previously published set of ARF5-inducible genes from seedlings^66^ (Figure S6E) indicating that we indeed revealed ARF5-dependent genes in stems. Strikingly, while our expectation was that genes, which are induced by *GR-ARF5ΔIII/IV* induction would be repressed by the auxin signaling repressor *bdl* and vice versa, we observed 144 genes (24%) that were either induced (73 genes) or repressed (71 genes) by both constructs (Figure 5A). This indicated that in PXY-positive cells ARF5 antagonizes the effect of overall auxin signaling on a substantial subset of target genes. Since we observed opposing effects of ARF5 and total canonical auxin signaling on cambium activity, we suspected that genes integrating these effects are among the 144 genes behaving in an unexpected manner. Interestingly, 11 genes out of 144 were also found in a set of genes that are differentially expressed during IC formation^57^ one of them being *WOX4* which was repressed by both GR-ARF5ΔIII/IV and Myc-GR-bdl induction (Figure 5B).

**Figure 5:**
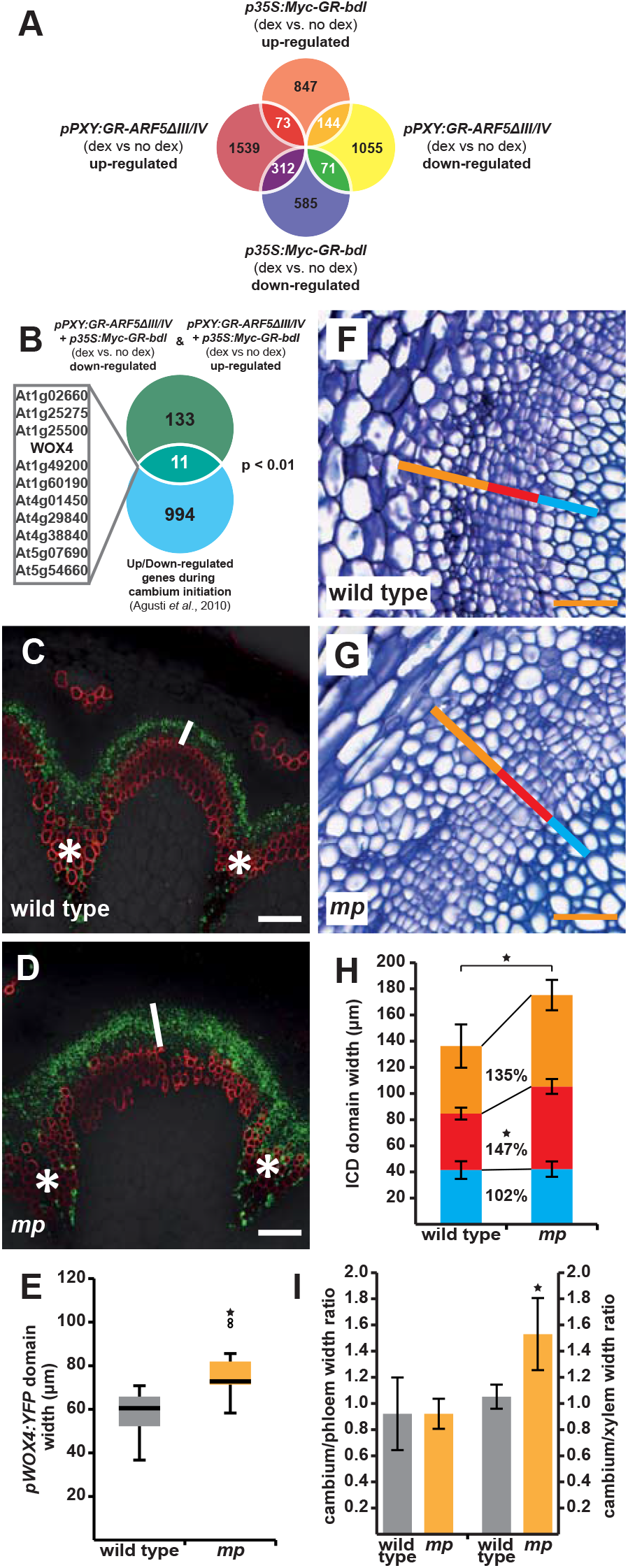
*A:* Venn-diagram of RNA sequencing results from RNA obtained from second internodes of *p35S:Myc-GR-bdl* and *pPXY-GR-ARF5ΔIII/IV* plants after mock or dexamethasone treatment. Identifiers of 600 overlapping genes are shown in Table S2. **B:** Comparison of the 144 unexpectedly acting genes and the group of genes up-or down-regulated during cambium initiation [57]. Non-random degree of the overlap was tested by using VirtualPlant 1.3 GeneSect with a cut-off p-value < 0.05 and the Arabidopsis thaliana Columbia (TAIR10) genome as background population. **C-D:** Confocal analyses of cross sections at the stem base of wild type and mp-S319 mutant plants containing pWOX4:YFP. White bars mark the *pWOX4:YFP* domain. Asterisks mark vascular bundles. Size bars represent 100 μm. **E:** Quantification of *pWOX4:YFP* domain width at the stem base of wild type and *mp-S319* plants. Student’s T-test was performed comparing wild type and *mp-S319* mutants (p=2.65E-03, sample size n=8-11). Significance is indicated by the asterisk. **F-G:** Toluidine blue stained cross sections at the stem base of wild type and *mp-S319* plants. Interfascicular regions are shown and the width of the vascular tissues phloem (orange bar), cambium (red bar) and xylem (blue bar) are marked. Size bars represent 50 μm. **H:** Quantification of the width of the vascular tissues phloem (orange), cambium (red) and xylem (blue) and the ICD at the stem base of wild type and *mp-S319* plants. Error bars: ± standard deviation. Student’s T-test was performed comparing wild type and mp-S319 mutants (phloem p=8.52E-02, cambium p=2.67E-04, xylem p=0.87 and ICD p=3.06E-02, Sample size n=5-6). Significance is indicated by the asterisk. **I:** Ratio of the radial extensions of cambium vs. phloem and cambium vs. xylem in wild type and *mp-S319* mutants. Student’s T-test (cambium/xylem p= 1.09E-02) or Welch’s T-test (cambium/phloem p=1.0) were performed comparing wild type and *mp-S319* mutants (Sample size n=5-6). Significance is indicated by the asterisk.

Because ARF5 induction resulted in both, *WOX4*-repression and the induction of xylem- and phloem-related genes (Figure S6), we reasoned that the repressive effect of ARF5 on cambium proliferation was due to an influence on the transition of cambial stem cells to vascular cells. To test this, we analyzed the stem cell marker *pWOX4:YFP* in *mp-S319* mutants. Indeed, the radial extension of the *pWOX4:YFP* domain was increased in *mp-S319* plants (Figure 5C-E) suggesting that the number of undifferentiated cambium cells was higher when *ARF5* activity was reduced. Consistently, when analyzing the anatomy of the cambium zone predominantly the size of the domain of undifferentiated cells was increased (Figure 5F-H) resulting specifically in an increased ratio of the domains of undifferentiated cells to xylem cells (Figure 5I). This indicated that ARF5 predominantly fulfils its function by promoting the transition of undifferentiated stem cells to differentiated xylem cells.

### *WOX4* mediates *ARF5* activity

Considering the role of *WOX4* as a mediator of auxin responses^23^ and its response to GR-ARF5ΔIII/IV induction, we hypothesized that ARF5 acts on cambium activity by regulating *WOX4.* Indeed, the expression domain of the transcriptional *pWOX4:YFP* reporter^23^ almost perfectly overlapped with *pARF5:mCherry* at the stem base (Figure 6A-C). Moreover, *WOX4* transcript levels were increased in *mp-S319* mutant stems (Figure 6D) demonstrating that the endogenous *ARF5* gene is required for the regulation of *WOX4.* Importantly, the negative effect of *GR-ARF5ΔIII/IV* induction on *WOX4* activity was also observed in the presence of the protein biosynthesis inhibitor cycloheximide (Cyclo) (Figure 6E), which was in line with a direct regulation of *WOX4* by ARF5. Consistently, transient expression of ARF5ΔIII/IV in cultured cells had, similar as on other genes directly repressed by ARF5^68^, a strong effect on the activity of a *pWOX4:LUC* promoter reporter, while this effect was only minor when ARF3 was expressed (Figure 6F). This suggested that, in comparison to ARF3, ARF5 substantially influenced the activity of the *WOX4* promoter. Indeed, neither cambium-specific nor global induction of *GR-ARF3* activity led to a significant change in *WOX4* expression in wild type or *arf3;arf4* double mutants although *IPT3,* a putative downstream target of *ARF3^69^,* was induced (Figure S7A-C). Furthermore, *WOX4* expression was not significantly altered in the *arf3;arf4* double mutant (Figure S7D) making it rather implausible that *ARF3* and *ARF4* act on cambium activity by regulating *WOX4.*

**Figure 6:**
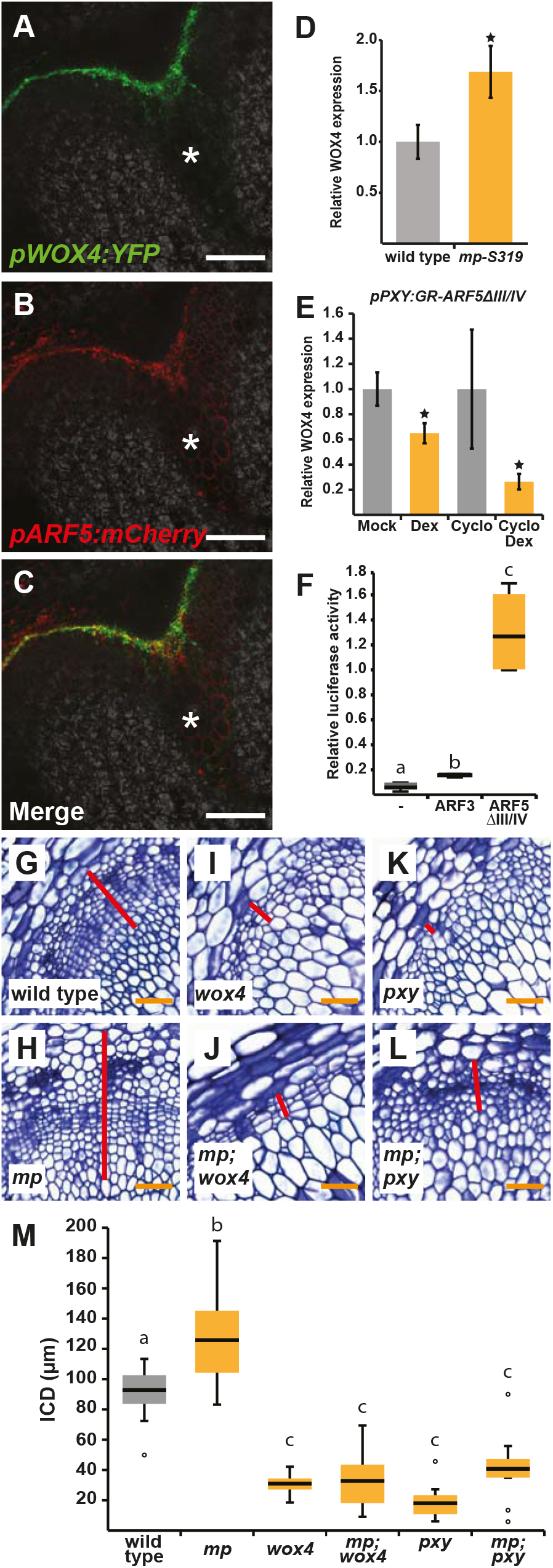
**A-C:** Confocal analysis of stem bases of plants carrying the *pWOX4:YFP* and the *pARF5:mCherry* reporter. Asterisks mark the vascular bundles. Size bars represent 100 μm. **D-E:** Analysis of *WOX4* transcript levels by quantitative RT-PCR at the stem base of wild type and *mp-S319* mutant pants (D) and in the second internode of *pPXY:GR-ARF5ΔIII/IV* plants after mock or Dex and cycloheximide or cycloheximide/Dex (E) treatments. Student’s T-test (wild type and *mp-S319* p=1.74E-02 and Mock and Dex p=1.66E-02) or Welch’s T-test (Cyclo and Cyclo/Dex p=1.29E-02) were performed comparing wild type and *mp-S319* mutants, mock and Dex and Cyclo and Cyclo/Dex, respectively (Sample sizes n=3-6). Significance is indicated by the asterisk. **F:** Analysis of relative luciferase activity of a *pWOX4:LUC* (firefly);p35S:LUC (Renilla) reporter in *Arabidopsis* protoplasts in the presence of *p35S:ARF3, p35S:ARF5ΔIII/IV* or no effector construct. Relative luciferase activity was determined by dual-luciferase assays. Statistical groups indicated by letters were determined by one-way ANOVA with post hoc Tamhane-T2 (CI 95%, Sample size n=4-5). **G-L:** Toluidine blue stained cross sections at the stem base of wild type and single and multiple mutant plants. Genotypes are indicated. Interfascicular regions are shown and ICD tissues are marked (red bar). Size bars represent 50 μm. **M:** Quantification of ICD tissue extension at the stem base of plants shown in G-L. Statistical groups indicated by letters were determined by one-way ANOVA with post hoc Tamhane-T2 (CI 95%, Sample size n=9-12).

To analyze the relevance of the observed effect of ARF5 on *WOX4* activity we determined ICD extension in *mp-S319* and *wox4-1* single and double mutants. While *mp-S319* showed enhanced cambium activity (Figure 6G, H, M), cambium activity was similar in *wox4-1* single and in *mp-S319;wox4-1* double mutants (Figure 6I, J, M) suggesting that *WOX4* is required for an ARF5-dependent repression of cambium activity. In comparison, depletion of *ARF5* activity in *mp-S319;pxy-4* double mutants lead to a mild suppression of cambium defects observed in *pxy-4* single mutants (Figure 6K, L, M^57^), suggesting that the epistatic relationship between *WOX4* and *ARF5* is specific.

## Discussion

Similar to apical meristems, the regulation of the vascular cambium has been tightly associated with the plant hormone auxin for several decades^13, 17, 70^. However, spatial organization of functional signaling domains and the role of auxin signaling in controlling different aspects of cambium activity remained unknown. Here, we show that auxin signaling takes place in cambium stem cells and that this signaling is crucial for cambium activity. We also show that not only stem cell activity in general but also the balance between undifferentiated and stem cells depends on the auxin signaling machinery with *ARF5* fulfilling a rather specific and *WOX4*-dependent role in this respect. Thus, auxin-related signaling controls distinct aspects of cambium activity important for a dynamic tissue production and a complex growth process.

The concentration of IAA peaks in the center of the cambial zone in *Populus* and Pinus^13, 14, 15^ and transcriptional profilings indicated a spatial correlation of this peak with the expression of auxin signaling components^17, 71^. However, genes responding to auxin were rather expressed in developing xylem cells arguing that sites of intense auxin signaling and of downstream responses do not necessarily overlap^18^. Consistently, our analysis of the highly sensitive auxin response marker *pDR5revV2:YFP* revealed a moderate auxin response in PXY-positive stem cells and a higher response in differentiated vascular tissues. Importantly, the auxin response in the PXY-positive region is overall pivotal for cambium activity since its local repression resulted in reduced tissue production similar as found in *wox4* or *pxy* mutants defective for canonical regulators of stem cell activity^23, 28, 57^. This demonstrates that, in the cambium, auxin signaling promotes stem cell activity in a cell-autonomous manner. Interestingly, *ARF5* and auxin signaling acts upstream of *WOX5* in the context of RAM organization^24, 72^ but differentiation of distal root stem cells is promoted by *ARF10* and *ARF16^72,^* ^73^. In comparison, *ARF5* restricts the stem cell domain in the SAM by repressing stem cell-related features^5^. In the RAM and the SAM, *ARF5* expression is found next to the expression domains of their central regulators *WOX5* and *WUS,* respectively^5, 24, 40^, whereas it overlaps completely with the domain of *WOX4* expression in the cambium. Thus, a division of labor of different auxin signaling components is found in various plant meristems and recruitment of distinct factors and adaptation of expression domains seem to have happened during the evolution of those systems.

*ARF5* plays a major role in translating auxin accumulation into the establishment of procambium identity in embryos and leaf primordia (reviewed in ^74^). However, ARF5 is also tightly associated with xylem formation via its direct targets *TMO5* and *ATHB8^65,^* ^75, 76^. In fact, we found both genes and their targets *ACAULIS5 (ACL5), SUPPRESSOR OF ACAULIS5 LIKE3 (SACL3*) and *BUSHY AND DWARF2* (BUD2)^21^, ^76^ to be induced upon *GR-ARF5ΔIII/IV* induction. Together with the observation that the domain of WOX4-positive cells is enlarged in *arf5* mutants, this suggests that *ARF5* promotes the transition of stem cells to xylem cells. Because we also revealed a negative effect of *ARF5ΔIII/IV* induction on *WOX4* activity, as well as a responsiveness of the *WOX4* promoter in transient expression systems and an epistatic genetic relationship between *WOX4* and *ARF5,* we propose that ARF5 fulfils its function partly by attenuating *WOX4* activity. Therefore, ARF5 acts as one hub modulating the activity of a multitude of genes in PXY-positive cells to foster the transition from stem cells to differentiated vascular cells. Consistent with the possibility that ARF5 does not necessarily act as a transcriptional activator, it represses *AUXIN RESPONSE REGULATOR15 (ARR15*) and *STOMAGEN* in the SAM and in leaf mesophyll cells, respectively^5, 68^. As in our case observed for *WOX4,* both genes are yet induced by ARF5 in transient expression systems^68^. The role of ARF5 in transcriptional regulation does therefore depend on the target promoter and the respective cellular environment^77^. Remarkably, not only xylem-related but also phloem-related genes are activated in stems when inducing *GR-ARF5ΔIII/IV* plants. This would argue for a general role of *ARF5* in promoting vascular differentiation and for the existence of one pool of stem cells marked by *PXY* promoter activity and feeding both xylem and phloem production. Alternatively, promotion of xylem differentiation is translated rapidly into the promotion of phloem differentiation by cell-to-cell signaling. The fact that the stem cell-to-phloem ratio is not altered in *arf5* mutants would argue for the latter option.

Consistent with a crucial role of cell-autonomous auxin signaling in cambium stem cells, *ARF3, ARF4* and *ARF5* expression was found in PXY-positive cells with *ARF5* being exclusively active in those. *ARF3* and *ARF4* have previously been shown to act in part redundantly in the establishment of abaxial identity in lateral organs^59^. In line with this function, we found both genes being mostly expressed distally of the cambium in phloem-related cell types. In fact, the lack of any effect on cambium or *WOX4* activity when modulating *ARF3* activity exclusively in PXY-positive cells, suggests that at least *ARF3* functions outside of this domain when regulating cambium activity and that ARF transcription factors positively regulating *WOX4* transcription still have to be discovered. Whether the phloem-related expression of *ARF3* and *ARF4* modulates the activity of cambium regulators expressed in areas distally to the *PXY* expression domain like MORE LATERAL GROWTH1 (MOL1)^29^ or CLAVATA/ESR41/44 (CLE41/44)^26, 27, 78^ remains to be determined. Transcriptionally, at least, our modulation of auxin signaling had no effect on the activity of *MOL1* or *CLE41/42* genes.

Collectively, we found a role of auxin signaling in the cambium sharing features with both the situation in the RAM where auxin regulates cell divisions^79^ and the SAM where auxin, and particular ARF5, is strongly correlated with cell differentiation^4^. Thereby, we enlighten a long-observed role of auxin signaling in plant development and reveal that its function is partly specific in different stem cell niches. The involvement of different auxin signaling components regulating individual aspects of meristem activity may provide a setup required for regulating a complex developmental process by one simple signaling molecule.

## Acknowledgements

This work was supported by the SFB 873 of the German Research Foundation (DFG) and a Heisenberg professorship to T.G. (DFG, GR2104/5-1). *arf4-2* mutants and the *pRPS5a:Myc-GR-bdl* line were kindly provided by Alexis Maizel (COS Heidelberg, Germany) and Gerd Jürgens (University of Tübingen, Germany), respectively. *mp-B4149* mutant seeds and the *pLC075* construct were obtained from Dolf Weijers (University of Wageningen, The Netherlands). Armin Djamei (GMI, Austria) donated the *pGreen-LUC-REN* construct. An established *Arabidopsis* (Col-0) dark-grown root cell suspension culture was a kind gift from Claudia Jonak (GMI, Austria). We thank members of the Greb lab for helpful discussions on the manuscript.

---

**Figure S1:**
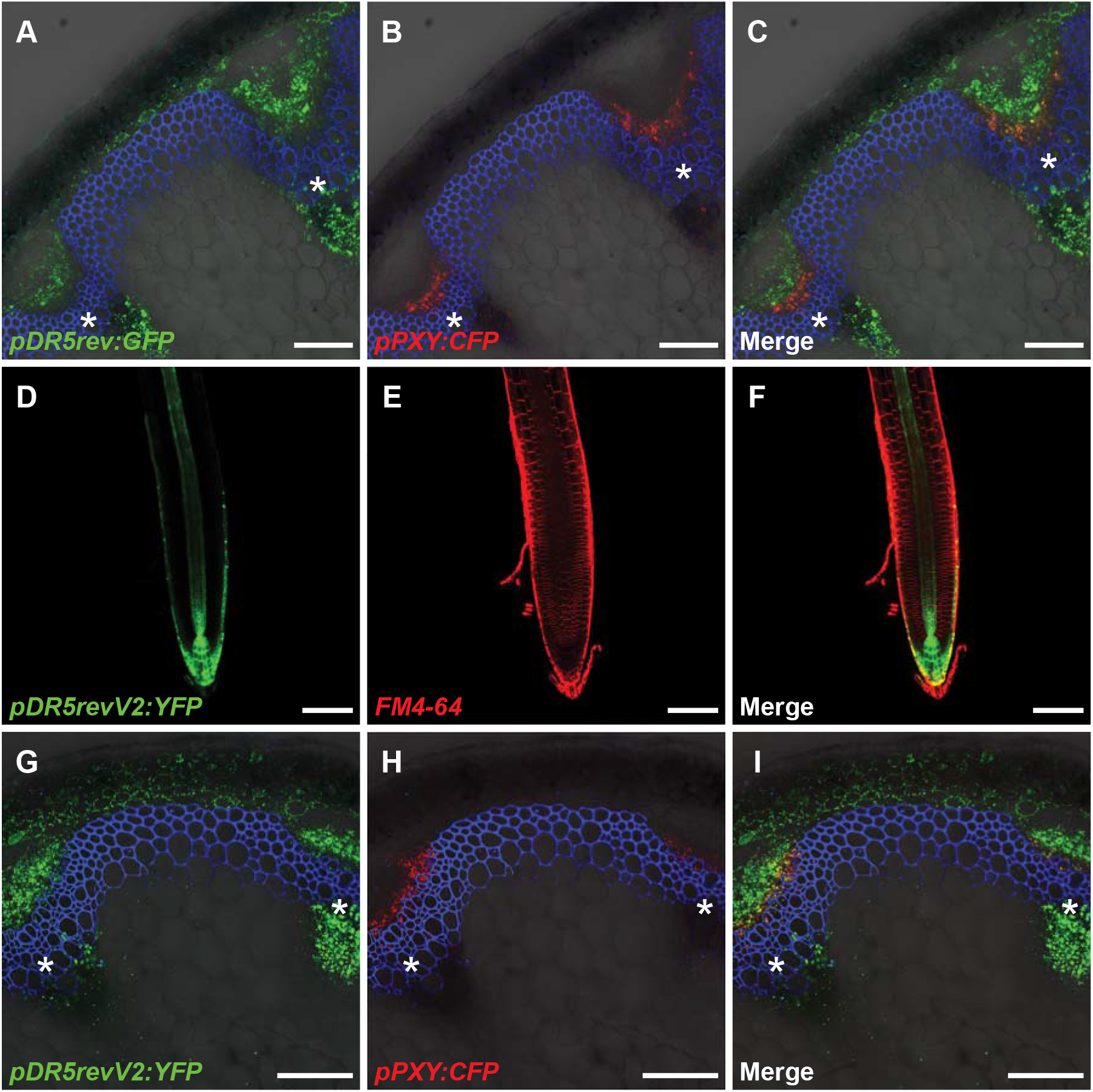
**A-C:** Confocal analyses of cross sections from the second internode (primary stem) of plants containing the auxin response markers *pDR5rev:GFP* and the stem cell marker *pPXY:CFP.* Asterisks mark the vascular bundles. Size bars represent 100 μm. PI staining in blue. **D-F:** Confocal analyses of root tips of 7-day-old seedlings carrying the *pDR5revV2:YFP* reporter. Size bars represent 100 μm. FM4-64 staining in red. **G-I:** Confocal analyses of cross sections from the second internode (primary stem) of plants containing the auxin response markers *pDR5revV2:YFP* and the stem cell marker *pPXY:CFP.* Asterisks mark the vascular bundles. Size bars represent 100 μm. PI staining in blue.

**Figure S2:**
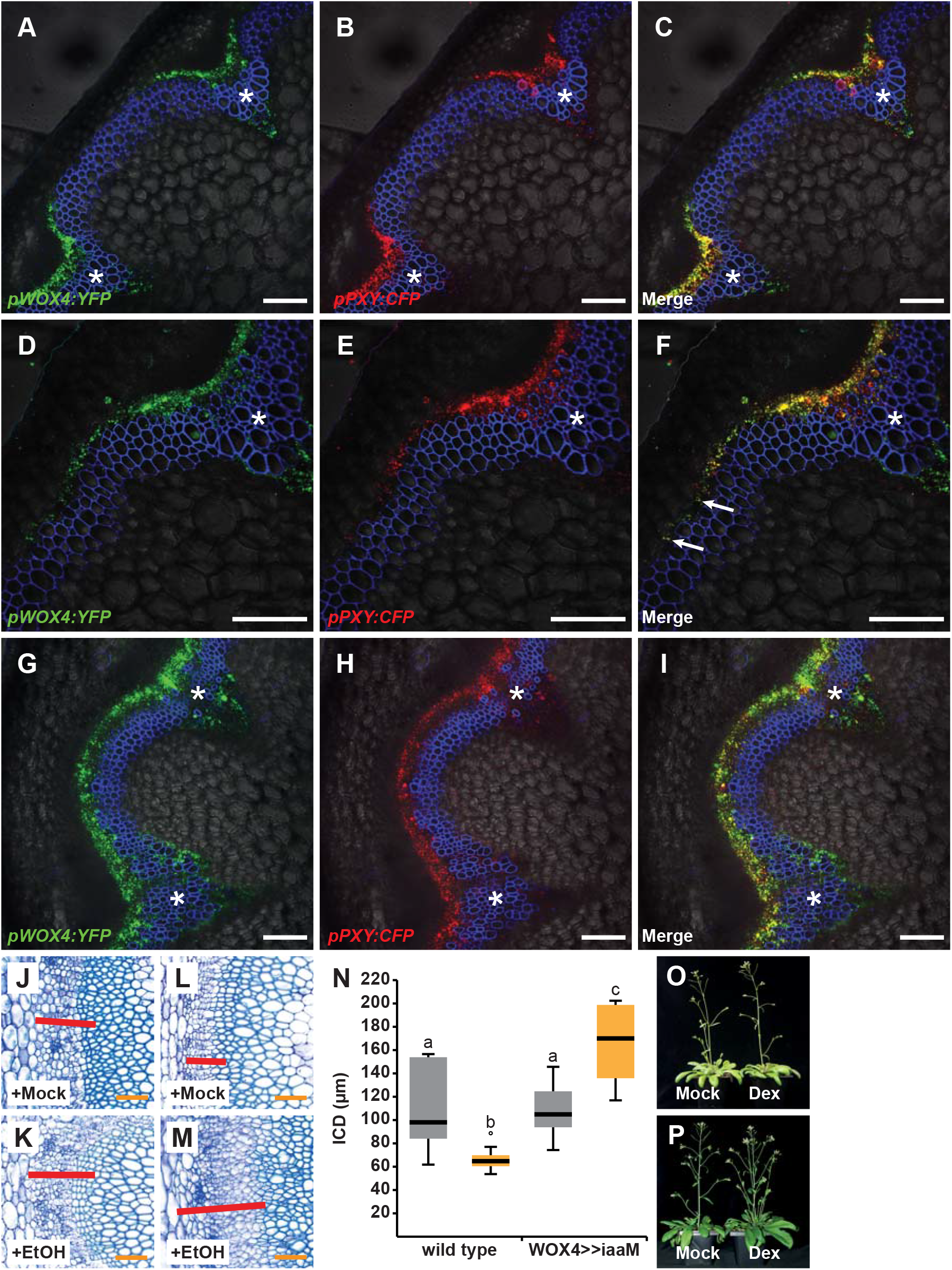
**A-I:** Confocal analyses of cross sections from the second internode (A-C), 2 cm above the stem base (D-F) and the stem base (G-I) of plants containing *pWOX4:YFP* and *pPXY:CFP.* Asterisks mark vascular bundles. Size bars represent 100 μm. PI staining in blue. **J-M:** Toluidine blue stained cross sections from the stem base of wild type (J, K) and *pWOX4:AlcR; pAlcA:iaaM* (L, M), plants after long-term EtOH (K, M) or mock (J, L) treatment. Interfascicular regions are shown and interfascicular cambium-derived tissues (ICD) are marked (red bar). Size bars represent 50 μm. **N:** Quantification of ICD tissue extension at the stem base of wild type and *pWOX4:AlcR; palcA:iaaM (WOX4>>iaaM*) plants after long-term EtOH (yellow), mock (light grey) treatment. Statistical groups indicated by letters were determined by one-way ANOVA with post hoc Tamhane-T2 (CI 95%, Sample size n=11-14). **O-P:** Overview pictures of 15-20 cm tall *pPXY:Myc-GR-bdl* (O) and *pBDL:Myc-GR-bdl* (P) plants after long-term mock or Dex treatments, respectively.

**Figure S3:**
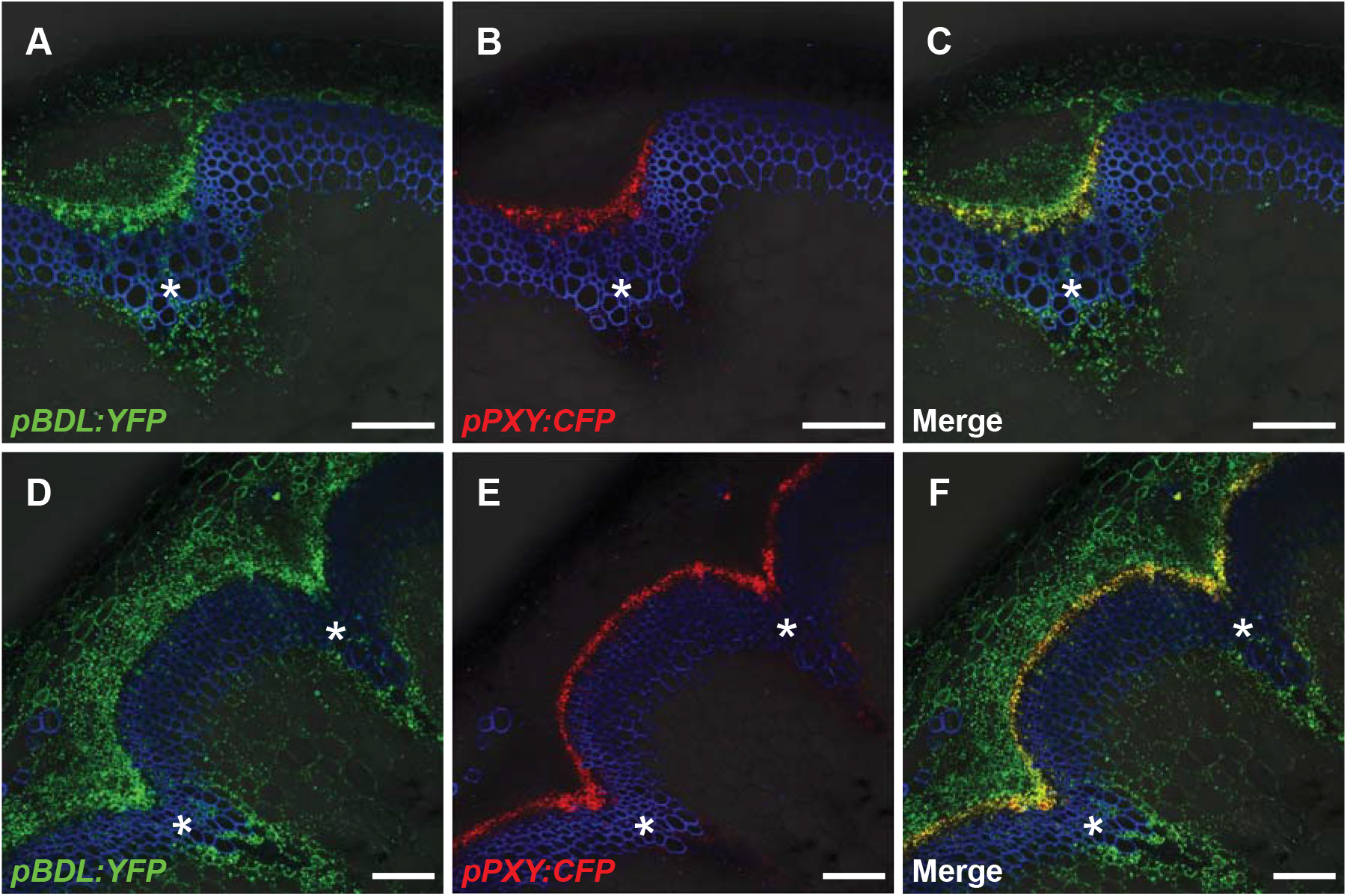
**A-F:** Confocal analyses of cross sections from the second internode (A-C) and stem base (D-F) of plants carrying *pBDL:YFP* and the stem cell marker *pPXY:CFP.* Asterisks mark vascular bundles. Size bars represent 100 μm. PI staining in blue.

**Figure S4:**
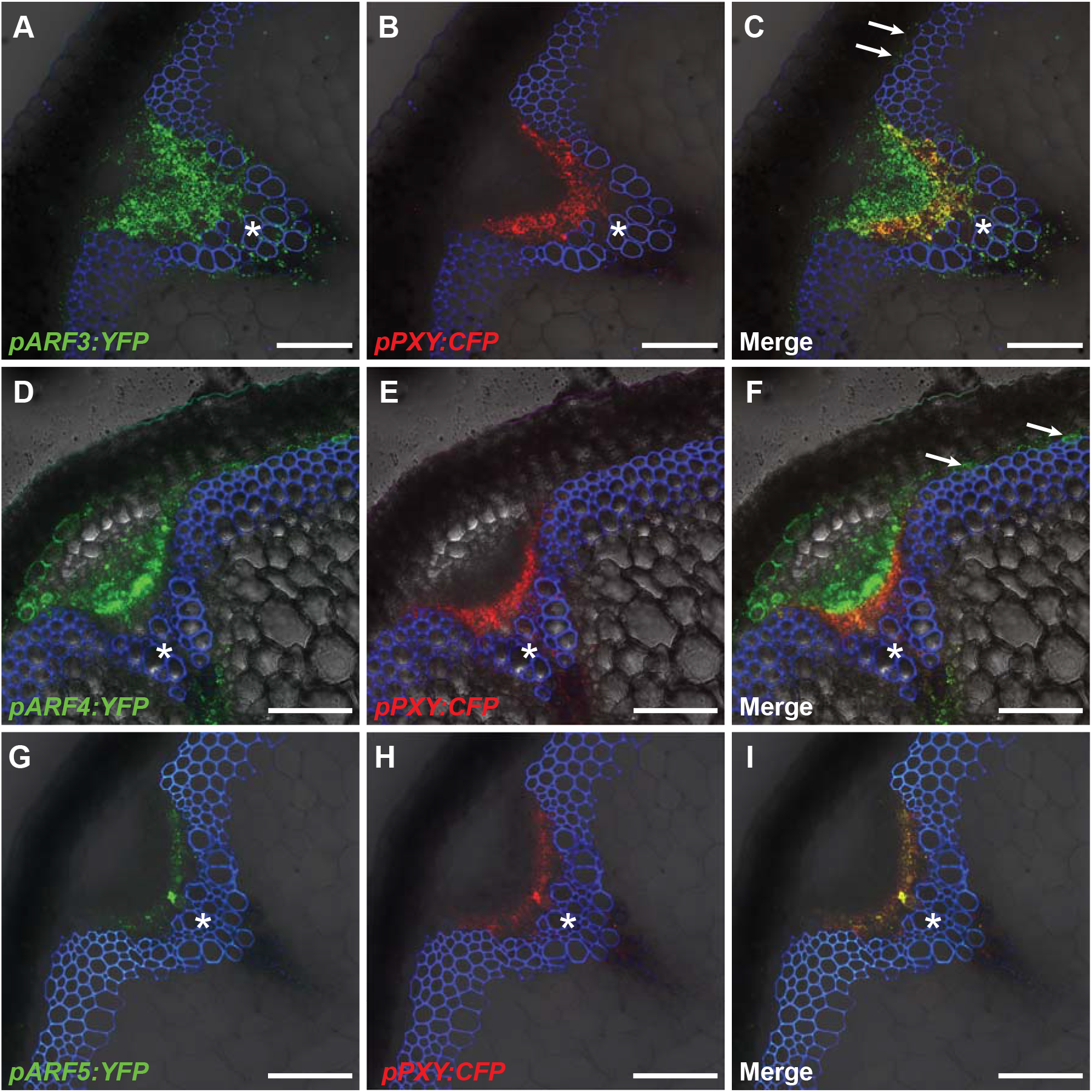
**A-I:** Confocal analyses of second internodes of plants containing *pARF3:YFP* (A-C), *pARF4:YFP* (D-F) and *pARF5:YFP* (G-I), respectively, and the stem cell marker *pPXY:CFP.* Arrows mark the expression of *pARF3:YFP* (C) and *pARF4:YFP* (F) in interfascicular region (starch sheath). Asterisks mark the vascular bundles. Scale bars represent 100 μm. PI staining in blue.

**Figure S5:**
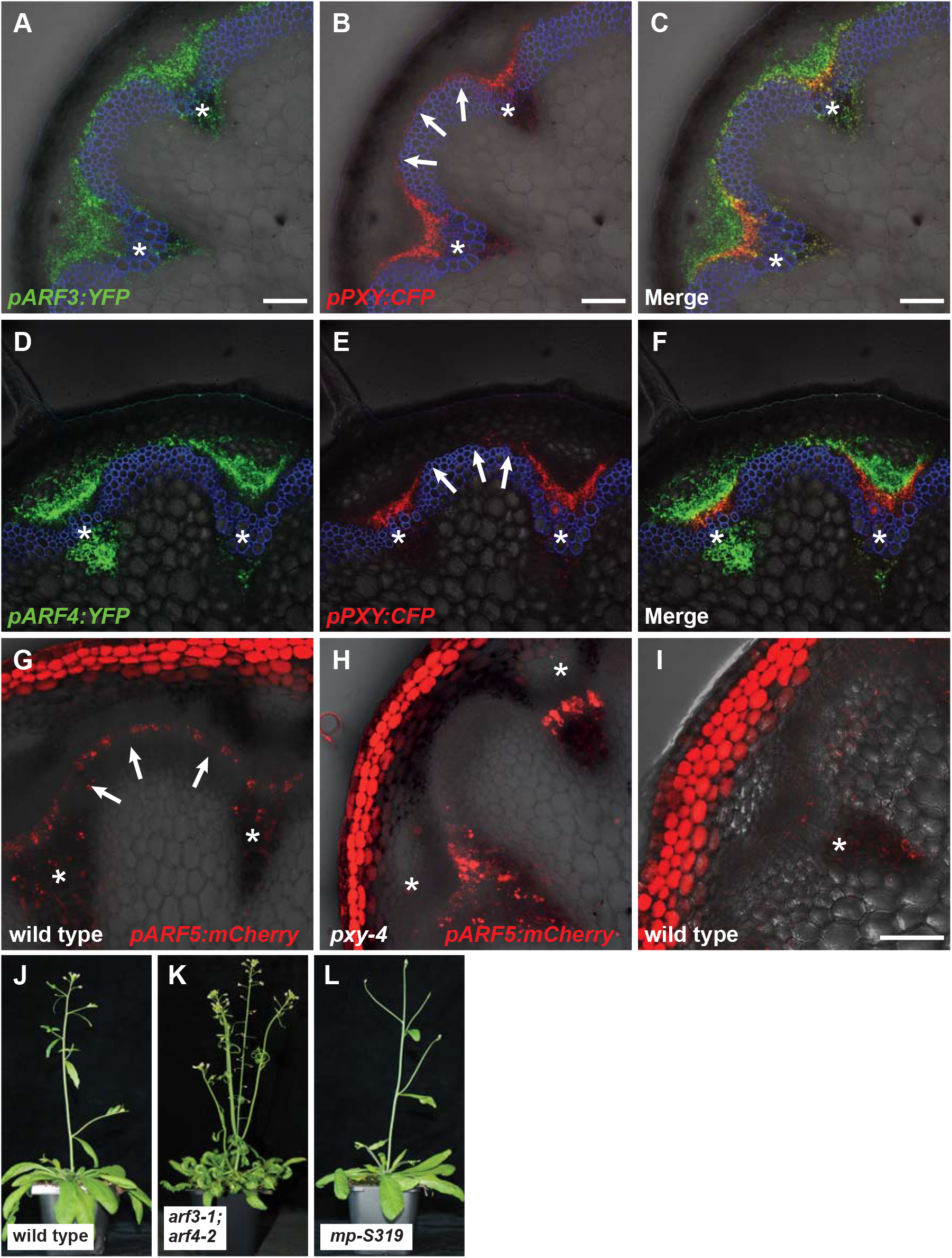
**A-F:** Confocal analyses of cross sections 5 mm above the stem base (transition zone) of plants containing *pARF3:YFP* (A-C) and *pARF4:YFP* (D-F), respectively, and the stem cell marker *pPXY:CFP.* Arrows indicate *pPXY:CFP* activity in interfascicular regions. Asterisks mark the vascular bundles. Size bars represent 100 μm. PI staining in blue. G-I: Confocal analyses of cross sections at the stem base from wild type and *pxy-4* mutant plants containing the *pARF5:mCherry* reporter. Asterisks mark the vascular bundles. Arrows indicate *pARF5:mCherry* activity in interfascicular regions. I shows auto-fluorescence at the stem base of wild type plants imaged with the same settings as in G and H. Size bars represent 100 μm. **J-L:** Growth habitus of wild type, *arf3-1;arf4-2* and *mp-S319* mutants.

**Figure S6:**
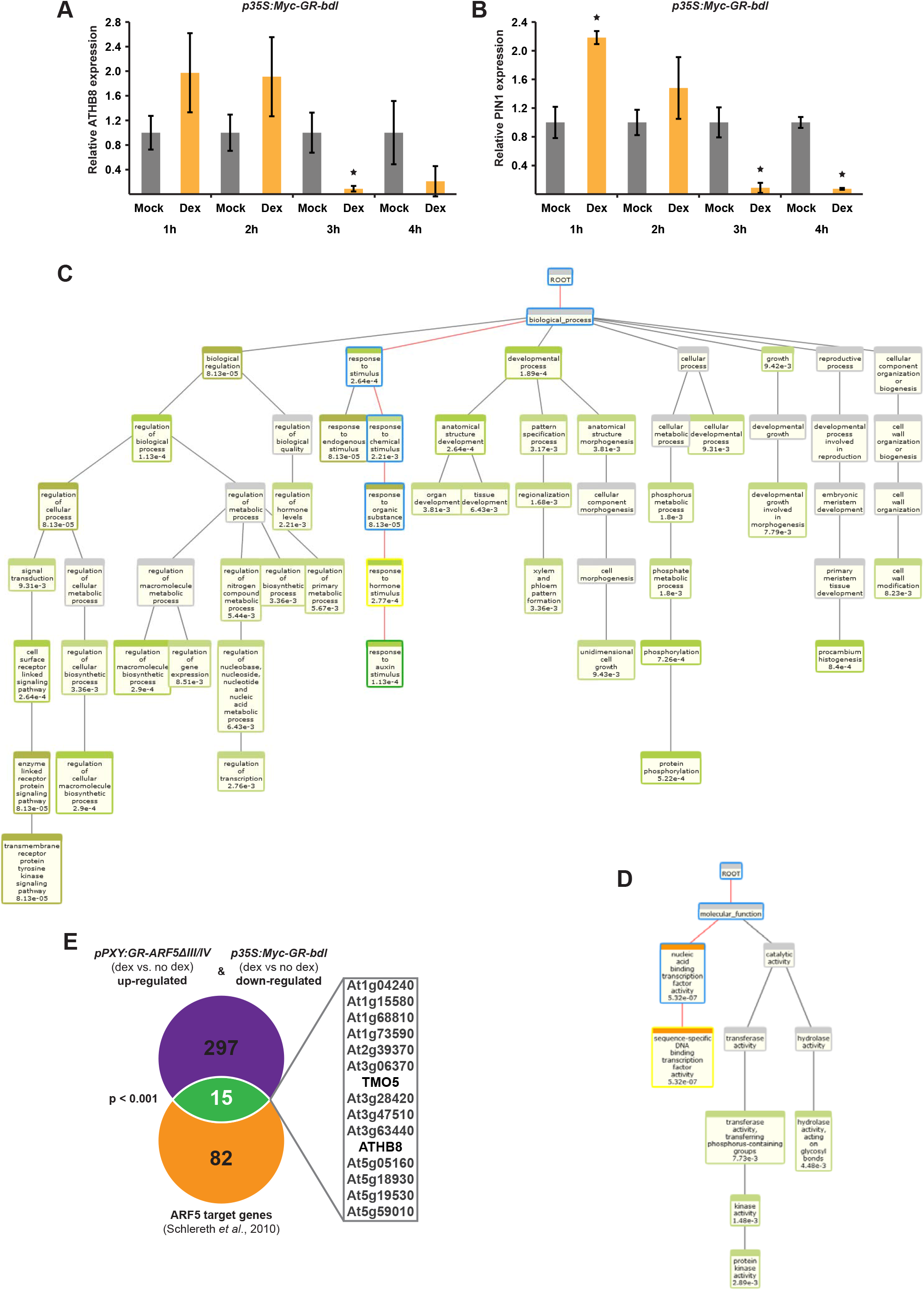
**A-B:** Optimization of the duration of Dex treatment in the second internode of *p35S:Myc-GR-bdl* plants. Shown are the analyses of transcript levels of *ATHB8* (A) or *PIN1* (B) by quantitative RT-PCR. Error bars: ± standard deviation. Student’s T-test (*ATHB8* 1h p=0.07, 2h p=0.11, 3h p=8.69E-03, 4 h p=0.19 and *PIN1* 1h p=9.86E-04, 2h p=0.06, 3h p0= 4.98E-03, 4H p=5.03E-04) was performed comparing mock and Dex treated samples (Sample sizes n=2-3). Significance is indicated by the asterisk. **C-D:** BioMaps analysis of biological process (C) and molecular function (D) of genes modulated by *p35S:Myc-GR-bdl* and *pPXY-GR-ARF5ΔIII/IV* induction (600 genes, p-value < 0.05, Table S2) by Virtual Plant 1.3 with a cut-off p-value < 0.01 and the *Arabidopsis thaliana* Columbia (TAIR10) genome as background population. **E:** Comparison of genes induced by *pPXY-GR-ARF5ΔIII/IV* and repressed by *p35S:Myc-GR-bdl* induction and a dataset of putative ARF5 target genes ^66^. Non-random degree of overlap was tested by using VirtualPlant 1.3 GeneSect with a cut-off p-value < 0.05 and the *Arabidopsis thaliana* Columbia (TAIR10) genome as background population.

**Figure S7:**
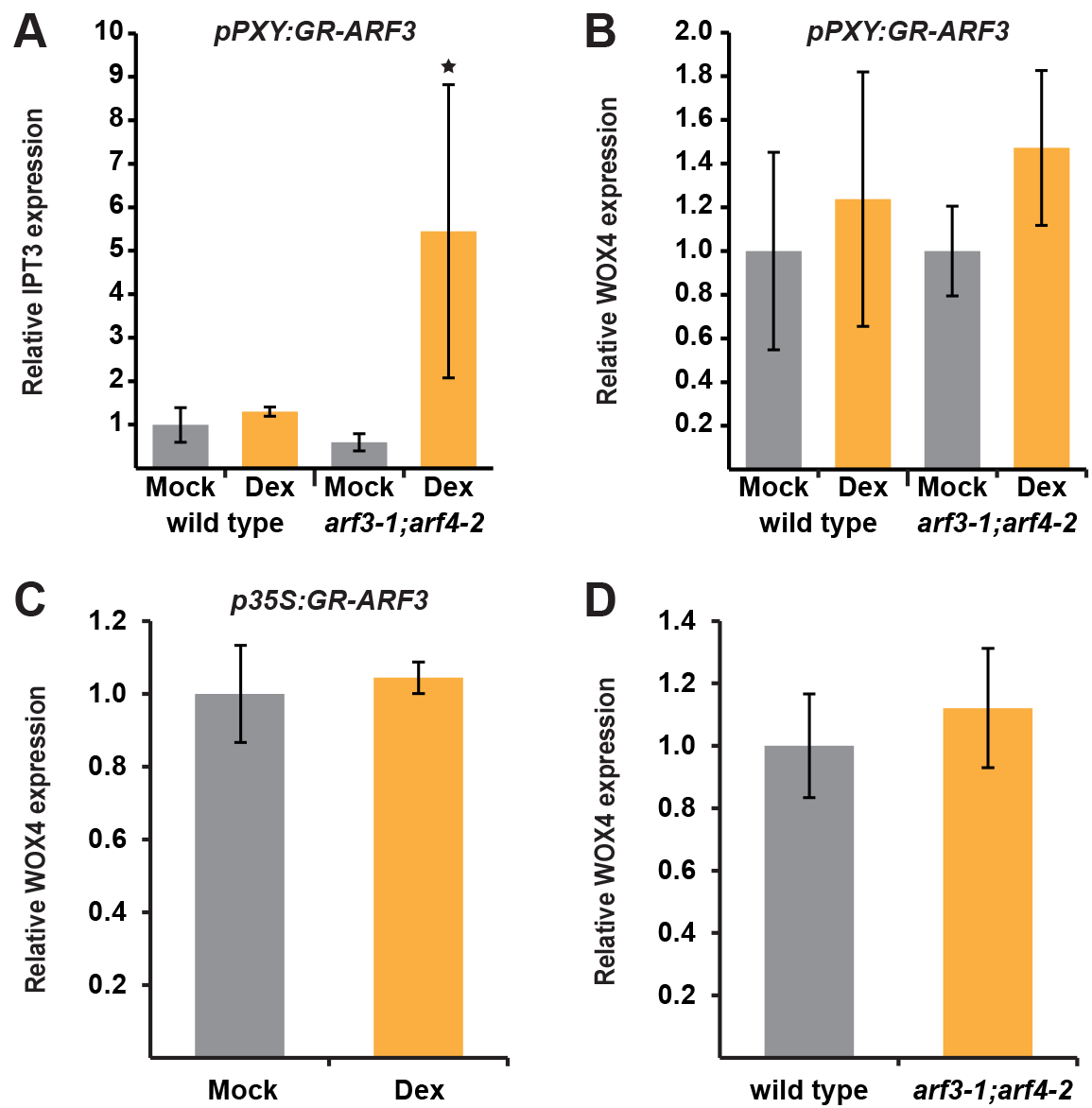
**A-B:** Analysis of *IPT3* (A) and *WOX4* (B) transcript levels by quantitative RT-PCR in the second internode of wild type and *arf3-1;arf4-2* mutant plants expressing *pPXY:GR-ARF3* after 3 hour mock (grey) or Dex (yellow) treatment. **C:** Analysis of *WOX4* transcript levels by quantitative RT-PCR in the second internode of wild type plants expressing *p35S:GR-ARF3* after 3 hour mock (grey) or Dex (yellow) treatment. **D:** Analysis of *WOX4* transcript levels by quantitative RT-PCR at the stem base of wild type and *arf3-1;arf4-2* mutant plants. Error bars: ± standard deviation. Student’s T-test (*pPXY:GR-ARF3 IPT3* p=0.17 & *WOX4* p=0.47, *pPXY:GR-ARF3;arf3-1;arf4-2 WOX4* p=0.17, *p35S:GR-ARF3 WOX4* p=0.61, wild type and *arf3-1;arf4-2 WOX4* p=0.45) or Welch’s T-test (*pPXY:GR-ARF3;arf3-1;arf4-2 IPT3* p=4.85E-02) were performed comparing mock and Dex and wild type and mutant, respectively (Sample size n=3-4). Significance is indicated by asterisk.

